# Posterior parietal cortex is causally involved in reward valuation but not probability weighting during risky choice

**DOI:** 10.1101/2023.02.08.527663

**Authors:** Ksenia Panidi, Alicia Nunez Vorobiova, Matteo Feurra, Vasily Klucharev

## Abstract

This study provides evidence that the posterior parietal cortex (PPC) is causally involved in risky decision making via the processing of reward values but not reward probabilities. In the within-group experimental design, participants performed a binary lottery choice task following transcranial magnetic stimulation of the right PPC, left PPC and a right PPC sham (placebo) stimulation. Both, mean-variance and the prospect theory approach to risky choice showed that the PPC stimulation changed participants’ preferences towards greater risk aversion compared to sham. On the behavioral level, after the PPC stimulation the likelihood of choosing a safer option became more sensitive to the difference in standard deviations between lotteries, compared to sham, indicating greater risk avoidance within the meanvariance framework. We also estimated the shift in prospect theory parameters of risk preferences after PPC stimulation. The hierarchical Bayesian approach showed moderate evidence (BF = 7.44 and 5.41 for right and left PPC respectively) for a credible change in risk aversion parameter towards lower marginal reward value (and, hence, lower risk tolerance), while no credible change in probability weighting was observed. Additionally, we observed anecdotal evidence (BF = 2.9) for a credible increase in the consistency of responses after the left PPC stimulation compared to sham.

## Introduction

The ability to make risky decisions is fundamental for survival. However, the brain mechanisms of risky behavior are still poorly understood. Previous studies have highlighted the role of the frontoparietal neural network as an important neural substrate for decision making under risk (Peter Mohr, Heekeren, & Rieskamp, 2017; Paulus et al., 2001). Several studies have shown that the dorsolateral prefrontal cortex (DLPFC) plays a fundamental role in risk taking (Huang et al., 2017; Knoch & Fehr, 2007; Pripfl, Neumann, Köhler, & Lamm, 2013; Ye et al., 2015). A meta-analysis reported that the activity in DLPFC is correlated with risk taking in the situation of active choice by contrast to merely anticipating the realization of a risky outcome (Mohr, Biele, & Heekeren, 2010). The role of the posterior parietal cortex (PPC) in making risky decisions has received much less attention in the neuroscientific literature.

Previous studies suggest that the PPC is involved in the assessment of risky options such as the degree of uncertainty of investments in the financial market (Peters & Büchel, 2009), with some subregions of the PPC decreasing activation in response to greater risk (van Duijvenvoorde et al., 2008). Making a risky decision is a complex process in which, as suggested by many economic studies, two components play a major role: subjective valuation of a monetary reward and probability weighting (Tversky & Kahneman, 1992). However, the neuroeconomic studies do not focus on the investigation of the role the PPC plays in these separate components. In the present study we aim to close this gap by causally addressing the PPC involvement in both subjective valuation of reward and probability weighting.

Previous studies allow us to hypothesize that PPC may be causally involved in both of these components. For example, a study on decisions under risk and ambiguity demonstrated that the PPC is more active in the situation of choice under risk rather than ambiguity in adolescents (Blankenstein, Schreuders, Peper, Crone, & van Duijvenvoorde, 2018). As the difference between risk and ambiguity consists in the availability of information about exact outcome probabilities, these findings support the hypothesis that the PPC might be involved in probability weighting. In a brain lesion study, participants with injuries in the PPC demonstrated impaired decision making in a risk-taking context compared to healthy controls, and the extent of the behavioral impairment correlated with the size of the lesion (Studer, Manes, Humphreys, Robbins, & Clark, 2015). Importantly, the perception of probabilities outside of decision-making contexts remained unimpaired. Another study revealed that repetitive transcranial magnetic stimulation (rTMS) of the temporoparietal junction shifted preferences towards lower risk taking particularly when outcome probabilities were 50%, suggesting the PPC might be also involved in risk sensitivity (Coutlee, Kiyonaga, Korb, Huettel, & Egner, 2016).

In the present study, we subjected participants to the three sessions of an offline rTMS protocol by using continuous theta burst stimulation (cTBS) over the right PPC, left PPC, and placebo stimulation over the right PPC, performed on separate days. In each session, after the stimulation participants made a series of binary lottery choices. The data allowed us to test the effects of TMS on risk taking using two different approaches to choice under risk currently existing in the literature. The first approach represents choice between lotteries as a mean-risk tradeoff (Markowitz, 1952; Tobin, 1958; Tobler et al., 2009). According to this approach, an individual gains positive utility from a higher expected value of a lottery, and obtains disutility from a higher variance (or standard deviation) of a lottery. The extent by which standard deviation affects the utility of a risky option is determined by an individual degree of risk tolerance (Grabenhorst et al., 2019). According to the second approach, known as prospect theory, an individual chooses between lotteries maximizing the sum of non-linear utilities of rewards weighted by non-linearly transformed probabilities (Tversky & Kahneman, 1992). As currently there is no consensus on which of these approaches better represents cognitive processes underlying risky choice (Boorman & Sallet, 2009; d’Acremont & Bossaerts, 2008; Dennison et al., 2022), we use both of them to analyze the data. Since the mean-variance approach implies that risk-sensitivity affects the choice linearly, we use this approach to analyze the data on a behavioral level with a linear regression model. Prospect theory, by contrast, depends on the curvature of utility and probability weighting functions Thus, we also use the structural modeling of risk preferences to determine the TMS effects on choice within a prospect theory framework.

On a behavioral level we find that both left and right PPC stimulation increased participants’ sensitivity to the difference in standard deviations between the riskier and the safer option (controlling for the difference in the mathematical expectations of the options), which corresponds to a decrease in risk tolerance within the mean-variance approach. The structural modeling of risk preferences estimated using the hierarchical Bayesian approach revealed that this shift in risk tolerance is associated with a credible decrease in the marginal value of money (risk aversion coefficient), while no such effect is found for probability weighting. Additionally, we observed that the TMS of the left PPC significantly decreased the amount of noise in participants’ choices.

## Materials and Methods

### Participants

We recruited 36 healthy volunteers (61% females, mean age = 22, min age = 19, max age = 27) who participated in all three sessions of the experiment. Participants were recruited via paper flyers distributed on the university campus as well as advertisements on the Internet. Potential subjects were queried about their area of education, and those with prior knowledge of economics or technical sciences (math, physics, computer science, etc.) were not invited to participate. These subjects were excluded due to possible knowledge of various theories of choice (e.g., expected utility, prospect theory) which might bias the outcomes — they may try to deliberately align their behavior with these theories or may engage in calculating mathematical expectation of lotteries.

Other exclusion criteria included regular sleep of less than 6 hours per day, self-reported left-handedness, history of brain injury or head trauma, being diagnosed with any psychiatric or neurological illness including epilepsy and migraines, family history of epilepsy, taking any prescription medication, and having metal objects inside the body. All participants read and signed the informed consent form prior to the experiment. All procedures were approved by the ethics committee of HSE university. One participant was excluded from data analysis due to misconception regarding the unlimited response time in the task. Overall, 35 participants were included in the final data set.

### Experimental task and payment

The experimental protocol closely follows the one used in a previous TMS study on the role of the DLPFC in risky choice (Panidi et al., 2022). The experimental task consisted of 85 self-paced binary lottery choice questions. Each question involved a choice between option A and option B, where each option represented a lottery (for a similar task, see (Holt & Laury, 2002)).

**Figure 1** presents an example of a screen subjects would see during the experiment. A participant had to indicate which lottery they would prefer to play by pressing one of two buttons on the keyboard located in front of them (indifference between lotteries was not allowed). All lotteries were purely in the gain domain. The monetary outcomes were presented in monetary units (MU) corresponding to the local currency with 1:1 conversion ratio. Participants were paid a 500 monetary unit (MU) participation fee (~22USD based on the BigMac index at the time of data collection) for each session and were informed that all payments would be administered at the very end of the third session. Additionally, they were informed that one answer from each session would be selected randomly and the lottery that was preferred in this particular question would be played out for real to determine the final payment the participant would receive for each session. Participants were informed about the outcome of each session only at the end of the third session to avoid interaction of this information with risky behavior. All the information regarding participation fee and the payment of additional rewards based on the choices in the experimental task was thoroughly conveyed to the participants in the instructions prior to the beginning of the task. On average, participants earned 934 MU (~41USD based on the BigMac index at the time of data collection) for the lotteries in three sessions in addition to the participation fee. Therefore, the amounts used in the task were meaningful for participants. Across all sessions, completion of the task took 7.8 minutes on average.

**Figure 1.**
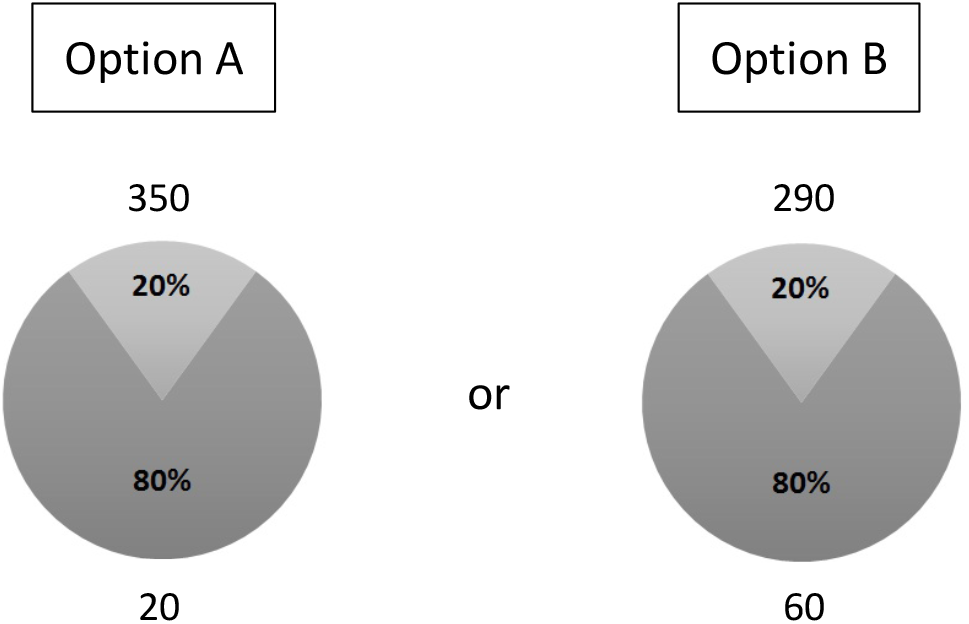
Task design. Subjects had to indicate their preferred option by pressing one of two buttons on the keyboard. The diagrams graphically and numerically present probability distributions for each lottery as well as the corresponding lottery outcomes.

In each trial, the lottery choice question was randomly taken from the list of 85 binary lottery choice questions. This list of questions consisted of five multiple price lists (MPL) similar to those used in other studies to detect changes in risk preferences (Holt & Laury, 2002). Each MPL represented a set of 17 binary choice questions ordered by probability of the best outcome ranging from 0 to 1. An example of an MPL used in the task is seen in **Table**. The complete list of lotteries used can be found in the Supplementary Materials.

In each pair of lotteries, the probabilities of the corresponding high and low outcomes were identical. The ordering of lotteries on the screen was randomized: in half of the questions the lottery with a higher outcome spread appeared as option A, while in the other half it appeared as option B. To minimize the possibility that subjects would remember their answers from previous sessions, the order of the questions was randomized and unique in each session. The positions of the monitor and keyboard were adjusted to suit each participant prior to the beginning of the task. To eliminate possible effects of time pressure on risk preferences found in previous studies (Kirchler et al., 2017), subjects were told that they had unlimited time for the task.

### Experimental design and stimulation protocol

For each participant, the experiment consisted of three sessions carried out on different dates separated by 3 to 4 days. In each session, we used a neuronavigated continuous theta-burst stimulation (cTBS) protocol. cTBS is an advanced patterned TMS protocol which has been shown to be effective in modulating the cortical excitability of a specific brain area both in motor and cognitive domains (Cho et al., 2010; Christov-Moore et al., 2016; Huang et al., 2005; Klucharev et al., 2011; Ott et al., 2011; Zack & Boileau, 2016). Each session included one of the three treatments: (1) cTBS of the right PPC (“right”), (2) cTBS of the left PPC (“left”), (3) sham stimulation of the right PPC (“sham”). The order of these treatments was randomized and counterbalanced between participants. To improve precision when positioning the coil, we employed a neuronavigation system which utilized the structural T1-weighted MRI scans which subjects obtained on a separate day prior to the experiment.

The stimulation was performed using a figure-of-eight (C-B60, 75mm diameter) coil through a MagVenture stimulator (MAGPRO R30 with MagOption, MagVenture, Inc.). The off-line stimulation paradigm was used; that is, stimulation was administered prior to performing the task. Stimulation intensity was set at 80% of the resting motor threshold (RMT) determined for each individual at the beginning of each session. The RMT was determined as the stimulation intensity inducing at least five motor evoked potentials (MEPs) of at least 50μV out of 10 pulses on the motor hotspot of the first dorsal interosseous muscle (Rossi et al., 2009) in the hand contralateral to the side of PPC stimulation. The cTBS stimulation lasted 40 seconds. The coil was held tangentially to the scalp at a 45-degree angle to the midsagittal axis of the subject’s head. Subjects were given a 5-minute break after the stimulation and before performing the task to allow for the downregulating effects of cTBS to take place (Huang et al., 2005). Previous research has shown that this stimulation protocol downregulates the cortex for up to 60 minutes following stimulation (Huang et al., 2005). Sham sessions were held in exactly the same way except that the stimulation was performed using a sham coil (MCF-P-65) which mimics the sound of the actual stimulation without inducing any perpendicular magnetic field to the cortex. Stimulation protocols were run with online neuronavigation (Localite GmbH, Germany).

Stimulation site coordinates were identified for each subject based on their T1-weighted structural MRI images. Montreal Neurological Institute (MNI) stereotaxic coordinates were back-normalized to subjects’ native brain space using an SPM8 toolbox (http://www.fil.ion.ucl.ac.uk/spm/software/spm8/). The stimulation coordinates (right PPC (42, −38, 44); left PPC (−36, −52, 46)) were selected based on a previous fMRI study (Peters & Büchel, 2009) where the peak activation at these sites correlated with the subjective value of a probabilistic reward. The right PPC coordinates are close to those reported in a meta-analysis (Mohr et al., 2010) where they were found to be more related to decision risk as opposed to anticipation risk. The left PPC coordinates are close to the coordinates used in a previous TMS study (Coutlee et al., 2016) which showed that downregulation of the left PPC in this area reduces risk taking behavior. TMS stimulation sites were identified on each participant’s scalp using the MRI-based Localite TMS Navigator system (Localite GmbH, Germany).

The level of discomfort in each session was assessed by means of self-report on the 7-point scale (1 indicating the lowest and 7 the highest experienced discomfort). The mean reported discomfort was as 1.5 for sham, 1.7 for right PPC and 1.9 for left PPC stimulation. The three sessions did not significantly differ in the reported discomfort (Wilcoxon signed-rank test p-value = 0.40 for sham vs. right PPC, 0.11 for sham vs. left PPC, 0.23 for left vs. right PPC).

### Behavioral analysis

To analyze the behavioral effects of TMS on risk taking we estimated a generalized linear mixed-effects regression model with a logit link function using the probability of choosing the riskier lottery as the dependent variable. In accordance with other studies on risk taking, we defined the riskier lottery as a lottery having higher standard deviation (Bougherara et al., 2021). We tested various model specifications with fixed effects for right and left PPC stimulation conditions, the difference in means in favor of the riskier lottery (Δ*μ*), the difference in standard deviations in favor of the riskier lottery (Δσ), the ratio Δ*μ*/Δ*σ*, and their interaction with stimulation conditions. Additionally, trial number and order of the experimental session were included to control for fatigue and order effects, as well as the selfreported level of discomfort during the stimulation to control for the emotional effects it might have had on participants’ decisions. Subject-level random effects were included in all model specifications.

The inclusion of the Δ*μ*/Δ*σ* ratio is suggested by the mean-risk approach to risky choice (Tobler, Christopoulos, O’Doherty, Dolan, & Schultz, 2009). The mean-risk approach postulates that participants derive utility from a higher mean of a lottery while experiencing disutility from a higher risk. The level of risk can be represented by the variance or the standard deviation of a lottery. The coefficient by which the risk measure is multiplied represents then the individual risk sensitivity. The utility of each lottery can then be represented as:

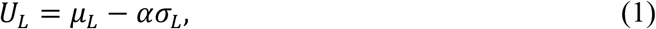

where *μ_L_* and *σ_L_* are the mean and standard deviation of a lottery respectively, and *α* > 0 is the degree of risk aversion. Following this framework, we can predict that a participant will choose the riskier lottery A versus the safer lottery B whenever *U_A_* > *U_B_*, i.e. whenever *μ_A_* – *μ_B_* > *α*(*σ_A_* – *σ_B_*). Therefore, in a deterministic model the participant will choose the riskier lottery whenever 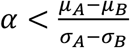. In the regression analysis, we expect a positive coefficient for the Δ*μ*/Δ*σ* regressor. A positive sign for this regressor would mean that participants are more likely to choose the riskier option when it offers a higher mean reward at the expense of a not very large difference in standard deviations with a safer option. Trials with probabilities equal to 0 or 1 were excluded from the analysis due to the standard deviation being zero and, as a result, Δ*μ*/Δ*σ* being undefined for these trials.

The advantage of the regression analysis is that it does not require specific assumptions regarding the functional forms for utility or probability weighting. However, the results might be biased since it does not allow for a non-linear relationship between various factors influencing risk-taking behavior. Therefore, we further proceed with the structural modeling of risk preferences and estimating the shift in preference parameters after the PPC stimulation.

### Structural modeling of risk preferences

To determine the effect of TMS stimulation on different components of risk preferences we estimated the structural model of choice under risk using the hierarchical Bayesian approach.

We assumed rank-dependent preferences with probability weighting in the form suggested in (Tversky & Kahneman, 1992), logistic distribution of the random error (Andersen, Harrison, Lau, & Rutström, 2008), a utility function with constant relative risk aversion (CRRA), and the difference in the logarithms of utility as a factor defining the choice between lotteries (strict utility) (Holt & Laury, 2002). The functional forms for probability weighting and utility are provided below:

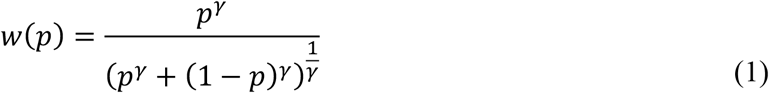

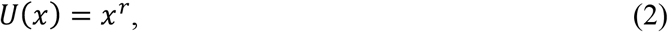

where *r* > 0 represents the risk-aversion coefficient, and *γ* represents the degree of probability distortion. The utility function is defined only in the gain domain since only positive outcomes were used in the experimental task. In accordance with the rank-dependent utility theory expected utility of lottery *L* is defined as 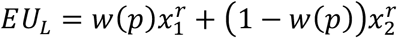, where *x*_1_ and *x*_2_ are the best and the worst monetary outcome respectively, and *p* is the probability of the best outcome. The probability of choosing lottery A over B is then given as:

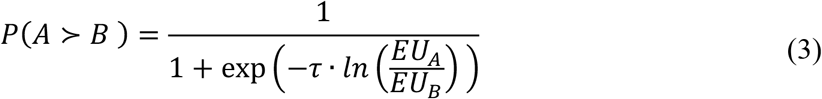

where *τ* represents the inverse “temperature”, or the inverse of the standard deviation of the noise, and *EU_A_* and *EU_B_* represent expected utilities of options A and B respectively.

Following the hierarchical Bayesian approach, we model each preference parameter as a combination of a baseline level and a change induced by the left or right PPC TMS stimulation:

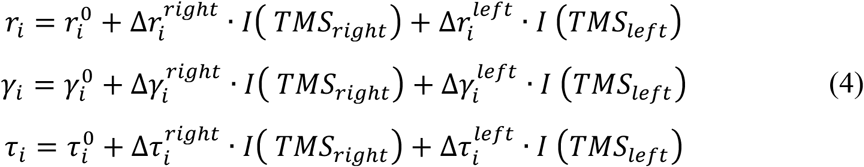

where *I*(·) equals 1 for trials from the corresponding TMS condition and 0 otherwise. The weakly informative priors for all unconstrained group-level parameters were taken from the standard normal distribution.

The standard deviations of all group parameters were sampled from a uniform distribution from 0 to 5. The individual-level parameters were linked to the unconstrained group-level parameters through Phi transformation, which would correspond to the uniform priors for constrained individual parameters. An additional linear transformation was applied to extend the support for these uniform distributions. We imposed the following restrictions on the individual and group parameter space: 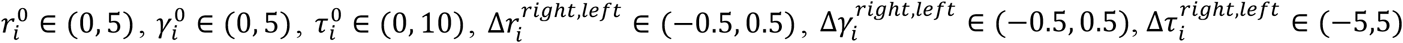. The obtained posterior distributions showed that all posterior samples well within these intervals and did not approach the boundaries.

The sampling was performed using the Markov Chain Monte Carlo (MCMC) method (NUTS algorithm) with 8 chains each containing 2,000 iterations for a warm-up and additional 2,000 iterations for sampling from posterior distribution giving 16,000 posterior samples for each parameter. Convergence was confirmed using the 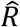 statistics and the visual inspection of the traceplots. The mean 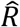 value for the group-level parameters equaled 1.009 with maximum value of 1.03, indicating that chains have mixed well. The estimates of model parameters were characterized with the mean of the posterior distribution for each group-level model parameter (denoted further as *μ*_*r*^0^_, *μ*_*γ*^0^_, *μ*_*τ*^0^_ for the baseline parameters and *μ*_Δ*r^left,right^*_, *μ*_Δ*γ^left,right^*_, *μ*_Δ*τ^left,right^*_ for the TMS effects for the left and right PPC respectively). The TMS effect was considered credible if the 95% Highest Density Interval for the corresponding variable did not contain zero. Additionally, for each parameter of interest we used the Savage-Dickey ratio at zero to compute the Bayes factor for testing the hypothesis that the group-level parameter is different from zero (i.e., testing *H*_1_ =: *δ* ≠ 0 against *H*_0_: *δ* = 0). We follow the standard interpretation of the Bayes factors with 1<BF<3 to indicate anecdotal, 3<BF<10 moderate, and BF>10 strong evidence in favor of the alternative hypothesis (Beard et al., 2016).

A posterior predictive check was performed by obtaining 4,000 random parameter samples from the joint posterior distribution. The proportion of correctly fitted choices was calculated for each selected sample to obtain the probability that the model fits participants’ choices correctly. This analysis indicated that the model fitted the participants’ choices correctly significantly better than chance (median = 0.80, 95% HDI= [0.79, 0.81]).

## Results

### Behavioral analysis

Table 2 presents the regression model estimation results. In all models, the dependent variable is the probability of choosing the riskier lottery in a trial. The regression analysis shows that regardless of the stimulation condition, participants behave in an expected way: they are more likely to choose the riskier lottery when it delivers, other things being equal, a higher mean reward compared to the safer one, when the difference in standard deviations with the safer lottery is smaller, and when the ratio Δ*μ*/Δ*σ* is higher.

**Table 1.** Example of the MPL used in the experimental task. Expected values were not shown on the screen during the task.

| Number of question | Option A |  | Option B |  | Probability of best outcome | Probability of worst outcome | Expected value of Option A | Expected value of Option B |
| --- | --- | --- | --- | --- | --- | --- | --- | --- |
|  | Outcome 1 | Outcome 2 | Outcome 3 | Outcome 4 |  |  |  |  |
| 1 | 750 | 15 | 450 | 250 | 0 | 1 | 15 | 250 |
| 2 | 750 | 15 | 450 | 250 | 0.01 | 0.99 | 22.35 | 252 |
| 3 | 750 | 15 | 450 | 250 | 0.05 | 0.95 | 51.75 | 260 |
| 4 | 750 | 15 | 450 | 250 | 0.10 | 0.90 | 88.5 | 270 |
| 5 | 750 | 15 | 450 | 250 | 0.15 | 0.85 | 125.25 | 280 |
| 6 | 750 | 15 | 450 | 250 | 0.20 | 0.80 | 162 | 290 |
| 7 | 750 | 15 | 450 | 250 | 0.30 | 0.70 | 235.5 | 310 |
| 8 | 750 | 15 | 450 | 250 | 0.40 | 0.60 | 309 | 330 |
| 9 | 750 | 15 | 450 | 250 | 0.50 | 0.50 | 382.5 | 350 |
| 10 | 750 | 15 | 450 | 250 | 0.60 | 0.40 | 456 | 370 |
| 11 | 750 | 15 | 450 | 250 | 0.70 | 0.30 | 529.5 | 390 |
| 12 | 750 | 15 | 450 | 250 | 0.80 | 0.20 | 603 | 410 |
| 13 | 750 | 15 | 450 | 250 | 0.85 | 0.15 | 639.75 | 420 |
| 14 | 750 | 15 | 450 | 250 | 0.90 | 0.10 | 676.5 | 430 |
| 15 | 750 | 15 | 450 | 250 | 0.95 | 0.05 | 713.25 | 440 |
| 16 | 750 | 15 | 450 | 250 | 0.99 | 0.01 | 742.65 | 448 |
| 17 | 750 | 15 | 450 | 250 | 1 | 0 | 750 | 450 |

**Table 2.** Mean-variance regression model estimation results. Dependent variable in all regressions: probability of choosing the riskier lottery. Columns (1) – (4) refer to model specifications with different sets of control variables and interaction terms.

|  | *Dependent variable:* |  |  |  |
| --- | --- | --- | --- | --- |
|  | Choose riskier lottery |  |  |  |
|  | (1) | (2) | (3) | (4) |
| Constant | -0.5043 | -0.3126 | -0.3193 | -0.3323 |
| Right PPC stimulation | -0.0938 | -0.0930 | -0.1403 | -0.1039 |
| Left PPC stimulation | -0.0434 | -0.0246 | -0.1001 | -0.0907 |
| Diff. in means / Diff. in stdv. | 0.7337*** | 0.7352*** | 0.6344*** | 0.6197*** |
| Diff. in means | 0.0068*** | 0.0068*** | 0.0068*** | 0.0071*** |
| Diff. in stdv. | -0.0080*** | -0.0080*** | -0.0080*** | -0.0079*** |
| Trial |  | 0.0015 | 0.0015 | 0.0015 |
| Order |  | -0.1178** | -0.1182** | -0.1190** |
| Discomfort |  | -0.0155 | 0.0077 | 0.0071 |
| Right PPC stimulation x Diff. in means / Diff. in stdv. |  |  | 0.1307* | 0.1799* |
| Left PPC stimulation x Diff. in means / Diff. in stdv. |  |  | 0.1964** | 0.1935* |
| Right PPC stimulation x Diff. in means |  |  |  | -0.0009 |
| Left PPC stimulation x Diff. in means |  |  |  | 0.00003 |
| Right PPC stimulation x Diff. in stdv. |  |  |  | -0.0002 |
| Left PPC stimulation x Diff. in stdv. |  |  |  | -0.0001 |
| Observations | 7,875 | 7,875 | 7,875 | 7,875 |
| Log Likelihood | -2,869.4490 | -2,865.0160 | -2,860.2390 | -2,859.7590 |
| Akaike Inf. Crit. | 5,752.8970 | 5,750.0320 | 5,744.4780 | 5,751.5170 |
| Bayesian Inf. Crit. | 5,801.6980 | 5,819.7470 | 5,828.1350 | 5,863.0610 |

*Note:* \* p<0.05, \*\* p<0.01, \*\*\* p<0.001

All regression model specifications suggest that the PPC stimulation does not have a direct effect on the probability of choosing the riskier lottery. However, both left and right PPC stimulation affects lottery choices by significantly changing the sensitivity of participants to the Δμ/Δσ ratio compared to sham (interaction terms *“Right PPC stimulation x Diff. in means /Diff. in stdv.*” and *“Left PPC stimulation x Diff. in means /Diff. in stdv.*”). This result indicates that after the PPC stimulation participants became more sensitive to the increase in the difference of the standard deviations between risky and safe options provided that the difference in their means is kept constant. In other words, when the standard deviation of the riskier lottery increases leading to a lower Δ*μ*/Δ*σ* ratio participants become less likely than they were in the sham session to choose the riskier lottery.

Therefore, behavioral analysis suggests that stimulation of either left or right PPC leads to greater sensitivity to risk (more risk-averse behavior).

### Parameter estimation

Prospect theory parameter estimation showed that in the sham session participants were risk averse on a group level with the risk aversion coefficient estimate being *μ*_*r*^0^_ = 0.80 (95% HDI = [0.44, 1.20]), demonstrated substantial probability distortion with average *μ*_*γ*^0^_ = 1.88 (95% HDI = [0.88, 2.95]), and were consistent in their choices with *μ*_*τ*^0^_ = 8.38 (95% HDI = [6.88, 9.81]). The group-level estimated TMS effects on model parameters with their respective 95% HDIs and Bayes Factors are presented in Table 3. Figure 2 shows the sampled posterior distributions of the TMS effect parameters.

**Table 3.** Estimated TMS effects on risk preference parameters: means, 95% HDIs and Bayes Factors for the changes in model parameters based on the sampled posterior distributions. The Bayes Factors indicate evidence in favor of *H*_1_: *δ* ≠ 0 against *H*_0_: *δ* = 0.

| Parameter | Mean | 95% HDI | BF |
| --- | --- | --- | --- |
| *TMS effects for right PPC* |  |  |  |
| $\Delta$ risk aversion ($\mu_{\Delta r}^{right}$) | -0.201 | [-0.35; -0.06] | 7.44 |
| $\Delta$ prob.weighting ($\mu_{\Delta \gamma}^{right}$) | 0.043 | [-0.01; 0.09] | 0.24 |
| $\Delta$ consistency ($\mu_{\Delta \tau}^{right}$) | 0.595 | [-0.99; 2.19] | 0.24 |
| *TMS effects for left PPC* |  |  |  |
| $\Delta$ risk aversion ($\mu_{\Delta r}^{left}$) | -0.175 | [-0.31; -0.05] | 5.41 |
| $\Delta$ prob.weighting ($\mu_{\Delta \gamma}^{left}$) | 0.047 | [-0.002; 0.1] | 0.35 |
| $\Delta$ consistency ($\mu_{\Delta \tau}^{left}$) | 2.089 | [0.298; 4.13] | 2.9 |

**Figure 2.**
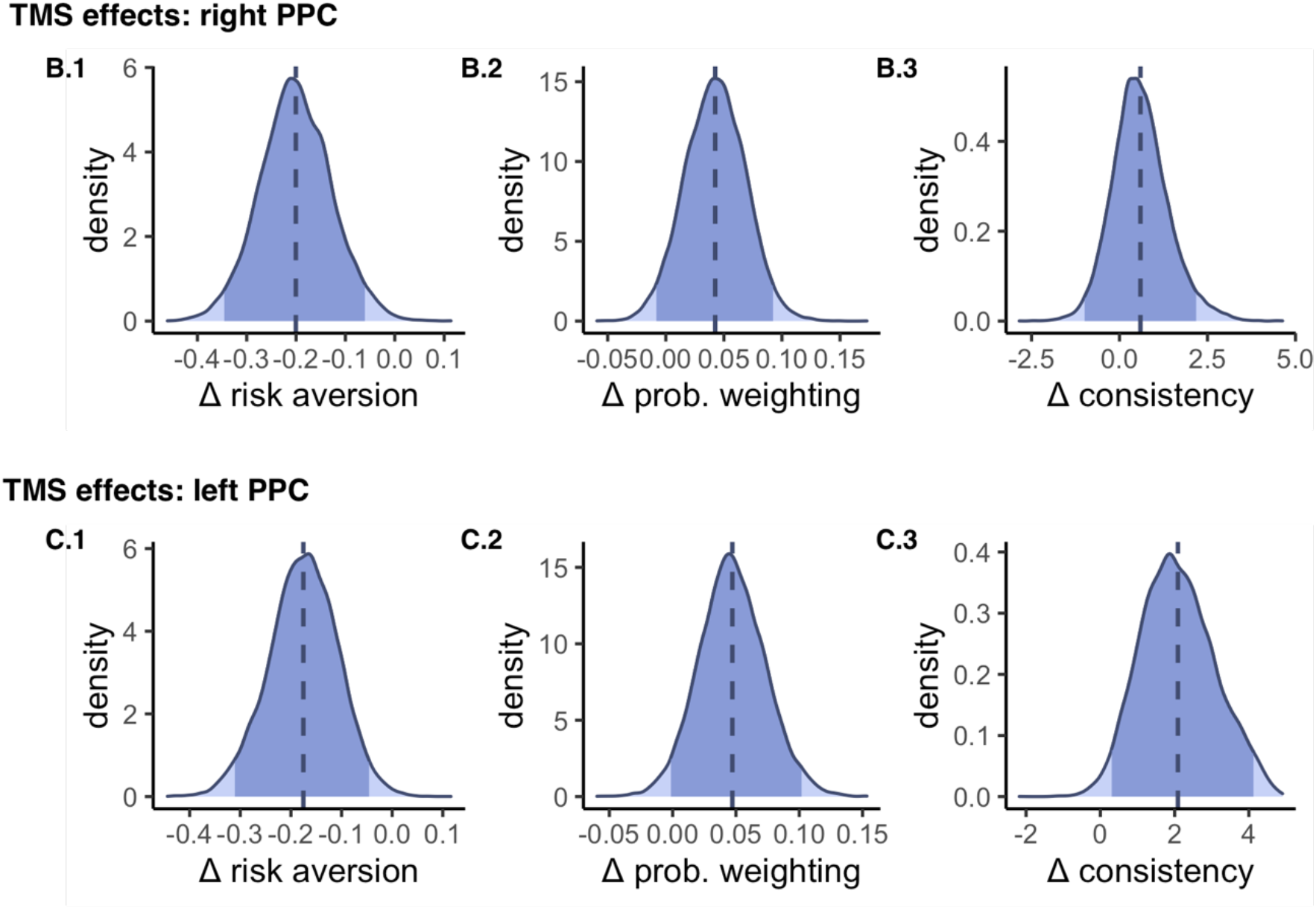
TMS effects on the risk preference parameters. Estimated posterior distributions are presented for: **B.1** change in risk aversion after right PPC TMS (*μ*_Δ*r^right^*_); **B.2** change in probability weighting after right PPC TMS (*μ*_Δ*γ^right^*_); **B.3** change in consistency after right PPC TMS (*μ*_Δ*τ^right^*_). **C.1** change in risk aversion after left PPC TMS (*μ*_Δ*r^left^*_); **C.2** change in probability weighting after left PPC TMS (*μ*_Δ*γ^left^*_); **C.3** change in consistency after left PPC TMS (*μ*_Δ*τ^left^*_). The shaded area under the curve corresponds to 95% HDI. Dashed vertical line indicates the mean (point estimate) of the posterior distribution. Zero is located outside the 95% HDIs for *μ*_Δ*r^right^*_ (figure B.1), *μ*_Δ*r^left^*_ (figure C.1), *μ*_Δ*τ^left^*_ (figure C.3) and indicating a significant change in the risk aversion parameter.

As can be seen from Table 3, the estimated HDIs for the change in risk aversion parameter after both left and right PPC stimulation do not contain zero, which suggests that the PPC stimulation leaded to a credible decrease in the marginal value of money (parameter *r* of the utility model). The Bayes factors indicate that the evidence for these effects is moderate. We do not observe any credible change in the probability weighting parameter. We do observe a positive shift in the consistency of preferences after the left PPC stimulation, however, the evidence for this effect is anecdotal as assessed by Bayes factor (self-reported discomfort did not significantly differ between the three stimulation conditions – see Methods).

## Discussion

The present study provides evidence that both left and right posterior parietal cortex is causally involved in risky decision making by its involvement in the processing of the marginal value of money rather than reward probability. In our experiment, downregulation of the PPC excitability leads to greater sensitivity to risk on a behavioral level relative to sham, as well as the downward shift in the estimated risk aversion parameter indicating decreased risk tolerance.

The results of the study are in line with the previously reported findings that the downregulation of the intraparietal sulcus with rTMS leads to a greater number of safe choices in trials with 50% probability lotteries (Coutlee et al., 2016). Our findings might give a clue as to why the shift in the number of safe choices was not previously observed for other probability levels (Coutlee et al., 2016): since a risky choice may depend on the valuation of reward and probability in a non-linear way, and given that participants may not always be consistent in their choices, the trial-by-trial analysis might not be enough to detect a significant shift in risk preferences. Therefore, structural modeling of risky choice might provide more information on the actual changes following the stimulation.

Further we discuss possible mechanisms which might have led to the observed effects of the PPC TMS on behavior. Importantly, these mechanisms might rely on the role of the PPC in representation of uncertainty, attention to salient stimuli, and numerical cognition.

Studies using animal models revealed that the PPC is involved in assessing the expectation of reward (Kobayashi, 2009). The lateral intraparietal (LIP) area activity in monkeys traced the desirability of an option based on the expected reward (Kubanek & Snyder, 2015), as well as the association of the rewarding option with a specific action needed to obtain it (Sugrue et al., 2004). It has also been shown that monkey parietal cortex neurons are strongly engaged in the saccades that reduce uncertainty (Horan et al., 2019). Single-cell recordings revealed that expected values and variances are encoded by separate populations of neurons in the fronto-parietal network, and that increased uncertainty enhances fronto-parietal bottom-up functional connectivity thereby increasing the amount of sensory input that would reduce the uncertainty (Taghizadeh et al., 2020). The results of the neuroanatomical studies suggest that a greater volume of gray matter in the PPC is correlated with higher tolerance to risk (Levy, 2017). These studies indicate that this brain area might play a role in stable personality-related components of risk preferences. Interestingly, the behavioral results of our study do not show the modulation of expected value effects on risky choice by TMS. Instead, we observe the increased sensitivity to the ratio between the expected value difference and the standard deviation difference in favor of the riskier option, which corresponds specifically to risk sensitivity.

Alternatively, the observed effects of the PPC stimulation might potentially be explained by the involvement of this region in numerical cognition (Roitman et al., 2012). In our task, the distance between the lottery outcomes might have been used by participants as a proxy for variance, or risk. It is well-known in numerical cognition that people more easily discriminate between the two numbers as the distance between them increases (Dehaene, 2007). If the TMS of the PPC temporarily affected the perception of difference between the monetary amounts, it might have shifted risk preferences as well. However, if numerical cognition was indeed distorted following the TMS stimulation, we would be likely to observe the distortion both in values and in probability perceptions since both modalities were numerically represented in the task. Since no credible changes were observed in the probability distortion parameter, we argue that the decrease in risk tolerance is not related to changes in numerical cognition but comes specifically from changes in the reward valuation process.

Importantly, the interpretation of the present findings crucially depends on the understanding of the underlying decision-making processes which guide participants’ choices between lotteries. Here, we based our analysis on the assumption that participants rely on multiplying and adding the weighted utilities of outcomes and aim to maximize the expected utility of an option. However, recent theories of choice under risk suggest that participants’ decisions may be based on various heuristic rules which simplify the decision-making process (Pachur et al., 2013). As the use of these heuristic rules may be reflected in the parameters of risk preferences (Pachur et al., 2017, 2018; Zilker & Pachur, 2021) the present study design does not allow to disentangle the use of heuristics and expected utility maximization.

Therefore, a plausible alternative explanation for the observed effects might be related to the modulation of attentional processes. In particular, TMS might have affected attention to salient rewards. A recent study in humans suggests that the value and salience of rewards are distinctly encoded in superior and inferior subregions of the PPC (Kahnt et al., 2014). Many previous studies also found that the activity in PPC is correlated with attention to information relevant for decision making (Corbetta & Shulman, 2002). Because stimulus salience is an important feature that guides attention, this finding opens the possibility that the PPC TMS might have altered participants’ visuo-spatial attentional processes during the accumulation of information about risky options. As a result, participants’ attention might have been shifted away from larger monetary amounts which are more salient but also more risky. The latter might have led them to prefer less risky options. The change in attentional processes might manifest itself in the TMS effects on the reaction time. To investigate this issue, we estimated a multilevel mixed effects linear regression model with the reaction time as a dependent variable and the stimulation conditions as the main variables of interest, including lottery characteristics, order of stimulation, trial number, and the self-reported level of discomfort as controls (see Supplemental material). Interestingly, we observed that after both left and right PPC stimulation the trial-by-trial reaction time significantly decreased compared to sham. At the same time, the change in reaction time might indicate that it became easier for participants to make a decision not to risk, for example, because they experienced less of a conflict between possible larger gain and chances to win only a small amount.

We also found anecdotal evidence (as assessed by the Bayes factor) that participants’ responses became less noisy after the left PPC stimulation. However, it is hard to interpret this finding directly since the understanding of the underlying nature of the inconsistencies in participants’ answers in this type of experimental tasks is largely lacking (Hollard et al., 2016). Economic experimental literature currently postulates two major sources of these inconsistencies. One hypothesis is that participants have stable and well-defined utility functions but they may make random errors at the moment of choice (‘random utility’ models). An alternative hypothesis is that participants’ preferences are not stable from trial to trial, i.e. the *parameters* of their utility function are subject to a random error in each trial (‘random preference’ models) (Loomes et al., 2002). In the present study, we applied random utility modeling of risk preferences which relies on the first interpretation of random errors. However, further analysis and fitting the data with random preferences rather than the random utility model may clarify the role of the PPC in the stability of risk preferences.

Several limitations of the study should be mentioned. First, to use a sufficiently large number of trials for a reliable estimation of the risk preference parameters we focused only on the gain domain. It has been suggested that gains and losses may be processed by separate neural networks in the brain (Mohr et al., 2010; Seymour et al., 2007; Zhang et al., 2018). Therefore, the conclusions of the present study shed light on the role of the PPC specifically in risky choices only when gains are involved. Second, in the stimulation procedure we focused on the specific coordinates in the PPC based on a previous fMRI study (Peters & Büchel, 2009).

However, the PPC is a large brain region with several subregions involved in various cognitive and motor functions. Hence, we cannot extrapolate the findings on the whole posterior parietal area. Finally, as a control condition we used sham stimulation only on the right PPC. This was done to reduce the repetition of the task for 4 times due to a within-subject design. Sham stimulation on the left PPC could also be used to control for the effects of discomfort on choice. Although the regression analysis did not reveal any significant effects of self-reported discomfort on the likelihood of choosing the riskier option, future studies are needed to control for left PPC placebo effects. Since in our study we used the offline stimulation protocol, our participants did not experience the discomfort of stimulation while performing the task itself, which could have biased the results if an online stimulation protocol had been used.

Overall, despite some limitations, our results demonstrate that both left and right PPC is causally involved in risky decision making, potentially via the marginal utility of reward. We do not find evidence for its involvement in probability weighting.

## Data and code availability

Complete dataset and custom code used to analyse the data are freely available in the repository: https://github.com/openaccessdata/PPC

## SUPPLEMENTAL MATERIALS

**Table S-1.** Complete list of lotteries used in the experimental task.

| Number of question | Option A |  | Option B |  | Probability of best outcome | Probability of worst outcome | Expected value of Option A | Expected value of Option B |
| --- | --- | --- | --- | --- | --- | --- | --- | --- |
|  | Outcome 1 | Outcome 2 | Outcome 3 | Outcome 4 |  |  |  |  |
| MPL 1 |  |  |  |  |  |  |  |  |
| 1 | 750 | 15 | 450 | 250 | 0 | 1 | 15 | 250 |
| 2 | 750 | 15 | 450 | 250 | 0.01 | 0.99 | 22.35 | 252 |
| 3 | 750 | 15 | 450 | 250 | 0.05 | 0.95 | 51.75 | 260 |
| 4 | 750 | 15 | 450 | 250 | 0.1 | 0.9 | 88.5 | 270 |
| 5 | 750 | 15 | 450 | 250 | 0.15 | 0.85 | 125.25 | 280 |
| 6 | 750 | 15 | 450 | 250 | 0.2 | 0.8 | 162 | 290 |
| 7 | 750 | 15 | 450 | 250 | 0.3 | 0.7 | 235.5 | 310 |
| 8 | 750 | 15 | 450 | 250 | 0.4 | 0.6 | 309 | 330 |
| 9 | 750 | 15 | 450 | 250 | 0.5 | 0.5 | 382.5 | 350 |
| 10 | 750 | 15 | 450 | 250 | 0.6 | 0.4 | 456 | 370 |
| 11 | 750 | 15 | 450 | 250 | 0.7 | 0.3 | 529.5 | 390 |
| 12 | 750 | 15 | 450 | 250 | 0.8 | 0.2 | 603 | 410 |
| 13 | 750 | 15 | 450 | 250 | 0.85 | 0.15 | 639.75 | 420 |
| 14 | 750 | 15 | 450 | 250 | 0.9 | 0.1 | 676.5 | 430 |
| 15 | 750 | 15 | 450 | 250 | 0.95 | 0.05 | 713.25 | 440 |
| 16 | 750 | 15 | 450 | 250 | 0.99 | 0.01 | 742.65 | 448 |
| 17 | 750 | 15 | 450 | 250 | 1 | 0 | 750 | 450 |
| MPL 2 |  |  |  |  |  |  |  |  |
| 1 | 480 | 105 | 400 | 180 | 0 | 1 | 105 | 180 |
| 2 | 480 | 105 | 400 | 180 | 0.01 | 0.99 | 108.75 | 182.2 |
| 3 | 480 | 105 | 400 | 180 | 0.05 | 0.95 | 123.75 | 191 |
| 4 | 480 | 105 | 400 | 180 | 0.1 | 0.9 | 142.5 | 202 |
| 5 | 480 | 105 | 400 | 180 | 0.15 | 0.85 | 161.25 | 213 |
| 6 | 480 | 105 | 400 | 180 | 0.2 | 0.8 | 180 | 224 |
| 7 | 480 | 105 | 400 | 180 | 0.3 | 0.7 | 217.5 | 246 |
| 8 | 480 | 105 | 400 | 180 | 0.4 | 0.6 | 255 | 268 |
| 9 | 480 | 105 | 400 | 180 | 0.5 | 0.5 | 292.5 | 290 |
| 10 | 480 | 105 | 400 | 180 | 0.6 | 0.4 | 330 | 312 |
| 11 | 480 | 105 | 400 | 180 | 0.7 | 0.3 | 367.5 | 334 |
| 12 | 480 | 105 | 400 | 180 | 0.8 | 0.2 | 405 | 356 |
| 13 | 480 | 105 | 400 | 180 | 0.85 | 0.15 | 423.75 | 367 |
| 14 | 480 | 105 | 400 | 180 | 0.9 | 0.1 | 442.5 | 378 |
| 15 | 480 | 105 | 400 | 180 | 0.95 | 0.05 | 461.25 | 389 |
| 16 | 480 | 105 | 400 | 180 | 0.99 | 0.01 | 476.25 | 397.8 |
| 17 | 480 | 105 | 400 | 180 | 1 | 0 | 480 | 400 |
| **MPL 3** |  |  |  |  |  |  |  |  |
| 1 | 350 | 20 | 290 | 60 | 0 | 1 | 20 | 60 |
| 2 | 350 | 20 | 290 | 60 | 0.01 | 0.99 | 23.3 | 62.3 |
| 3 | 350 | 20 | 290 | 60 | 0.05 | 0.95 | 36.5 | 71.5 |
| 4 | 350 | 20 | 290 | 60 | 0.1 | 0.9 | 53 | 83 |
| 5 | 350 | 20 | 290 | 60 | 0.15 | 0.85 | 69.5 | 94.5 |
| 6 | 350 | 20 | 290 | 60 | 0.2 | 0.8 | 86 | 106 |
| 7 | 350 | 20 | 290 | 60 | 0.3 | 0.7 | 119 | 129 |
| 8 | 350 | 20 | 290 | 60 | 0.4 | 0.6 | 152 | 152 |
| 9 | 350 | 20 | 290 | 60 | 0.5 | 0.5 | 185 | 175 |
| 10 | 350 | 20 | 290 | 60 | 0.6 | 0.4 | 218 | 198 |
| 11 | 350 | 20 | 290 | 60 | 0.7 | 0.3 | 251 | 221 |
| 12 | 350 | 20 | 290 | 60 | 0.8 | 0.2 | 284 | 244 |
| 13 | 350 | 20 | 290 | 60 | 0.85 | 0.15 | 300.5 | 255.5 |
| 14 | 350 | 20 | 290 | 60 | 0.9 | 0.1 | 317 | 267 |
| 15 | 350 | 20 | 290 | 60 | 0.95 | 0.05 | 333.5 | 278.5 |
| 16 | 350 | 20 | 290 | 60 | 0.99 | 0.01 | 346.7 | 287.7 |
| 17 | 350 | 20 | 290 | 60 | 1 | 0 | 350 | 290 |
| **MPL 4** |  |  |  |  |  |  |  |  |
| 1 | 680 | 70 | 430 | 380 | 0 | 1 | 70 | 380 |
| 2 | 680 | 70 | 430 | 380 | 0.01 | 0.99 | 76.1 | 380.5 |
| 3 | 680 | 70 | 430 | 380 | 0.05 | 0.95 | 100.5 | 382.5 |
| 4 | 680 | 70 | 430 | 380 | 0.1 | 0.9 | 131 | 385 |
| 5 | 680 | 70 | 430 | 380 | 0.15 | 0.85 | 161.5 | 387.5 |
| 6 | 680 | 70 | 430 | 380 | 0.2 | 0.8 | 192 | 390 |
| 7 | 680 | 70 | 430 | 380 | 0.3 | 0.7 | 253 | 395 |
| 8 | 680 | 70 | 430 | 380 | 0.4 | 0.6 | 314 | 400 |
| 9 | 680 | 70 | 430 | 380 | 0.5 | 0.5 | 375 | 405 |
| 10 | 680 | 70 | 430 | 380 | 0.6 | 0.4 | 436 | 410 |
| 11 | 680 | 70 | 430 | 380 | 0.7 | 0.3 | 497 | 415 |
| 12 | 680 | 70 | 430 | 380 | 0.8 | 0.2 | 558 | 420 |
| 13 | 680 | 70 | 430 | 380 | 0.85 | 0.15 | 588.5 | 422.5 |
| 14 | 680 | 70 | 430 | 380 | 0.9 | 0.1 | 619 | 425 |
| 15 | 680 | 70 | 430 | 380 | 0.95 | 0.05 | 649.5 | 427.5 |
| 16 | 680 | 70 | 430 | 380 | 0.99 | 0.01 | 673.9 | 429.5 |
| 17 | 680 | 70 | 430 | 380 | 1 | 0 | 680 | 430 |
| **MPL 5** |  |  |  |  |  |  |  |  |
| 1 | 335 | 10 | 230 | 110 | 0 | 1 | 10 | 110 |
| 2 | 335 | 10 | 230 | 110 | 0.01 | 0.99 | 13.25 | 111.2 |
| 3 | 335 | 10 | 230 | 110 | 0.05 | 0.95 | 26.25 | 116 |
| 4 | 335 | 10 | 230 | 110 | 0.1 | 0.9 | 42.5 | 122 |
| 5 | 335 | 10 | 230 | 110 | 0.15 | 0.85 | 58.75 | 128 |
| 6 | 335 | 10 | 230 | 110 | 0.2 | 0.8 | 75 | 134 |
| 7 | 335 | 10 | 230 | 110 | 0.3 | 0.7 | 107.5 | 146 |
| 8 | 335 | 10 | 230 | 110 | 0.4 | 0.6 | 140 | 158 |
| 9 | 335 | 10 | 230 | 110 | 0.5 | 0.5 | 172.5 | 170 |
| 10 | 335 | 10 | 230 | 110 | 0.6 | 0.4 | 205 | 182 |
| 11 | 335 | 10 | 230 | 110 | 0.7 | 0.3 | 237.5 | 194 |
| 12 | 335 | 10 | 230 | 110 | 0.8 | 0.2 | 270 | 206 |
| 13 | 335 | 10 | 230 | 110 | 0.85 | 0.15 | 286.25 | 212 |
| 14 | 335 | 10 | 230 | 110 | 0.9 | 0.1 | 302.5 | 218 |
| 15 | 335 | 10 | 230 | 110 | 0.95 | 0.05 | 318.75 | 224 |
| 16 | 335 | 10 | 230 | 110 | 0.99 | 0.01 | 331.75 | 228.8 |
| 17 | 335 | 10 | 230 | 110 | 1 | 0 | 335 | 230 |

**Table S-2.** Regression analysis of PPC TMS effects on reaction time.

|  | Dependent variable: |
| --- | --- |
|  | *Reaction time* |
| Constant | 8,983.30***<br>(637.17) |
| Right PPC stimulation | -760.30***<br>(144.20) |
| Left PPC stimulation | -355.25*<br>(147.75) |
| Probability of best outcome | 1,026.59*<br>(419.09) |
| Trial number | -20.87***<br>(2.38) |
| Difference in means | 2.17*<br>(0.84) |
| Difference in std.dev. | -0.33<br>(0.71) |
| (Diff. in means/ Diff. in std.dev.) | -162.21**<br>(54.95) |
| Discomfort | 467.47***<br>(86.12) |
| Session order | -1,618.61***<br>(73.90) |
| Observations | 7,875 |
| Log Likelihood | -78,565.10 |
| Akaike Inf. Crit. | 157,154.20 |
| Bayesian Inf. Crit. | 157,237.90 |
| *Note:* | * p<0.05 **p<0.01, ***p<0.001<br>Standard errors in parentheses. |

### Parameter recovery

For parameter recovery we simulated 729 each containing responses of 35 synthetic subjects with exactly the same lottery sets as in the real experiment. Since the model contains 9 group-level parameters, assigning 3 values to each of them and taking all possible permutations would imply 19,683 combinations. To reduce the number of simulated datasets we assigned only one value for each of the group-level baseline parameters. Each of the 6 parameters corresponding to the TMS effects was assigned 3 possible values. The complete permutation then generated 729 group-level parameter sets. Group-level parameters were chosen in the range similar to the values estimated on the real data as follows: *μ*_*r*^0^_ = 0.8, *μ*_*γ*^0^_ = 1.5, *μ*_*τ*^0^_ = 8, *μ*_Δ*r^right^*_ = (−0.3,-0.2,-0.1), *μ*_Δ*γ^right^*_ = (0.01,0.03,0.05), *μ*_Δ*τ^right^*_ = (−0.5,0.5,1), *μ*_Δ*r^left^*_ = (−0.25,-0.15,-0.05), *μ*_Δ*γ^left^*_ = (0.04,0.06,0.08), *μ*_Δ*τ^left^*_ = (0.5,1,2). The group-level standard deviations for the baseline parameters were selected as follows: *σ_r^0^_* = 0.1, *σ_γ^0^_* = 0.5, *σ_τ^0^_* = 0.1. Group-level standard deviations for the TMS effect parameters were set equal to those estimated on the real data.

For each group-level parameter value, the corresponding subject-level parameters were sampled from a uniform distribution. The support interval for the uniform distribution was selected to match the group-level parameter value as a mean and its corresponding standard deviation.

Each of 729 simulated datasets was then submitted to the same estimation pipeline as the real data. For each group-level parameter value estimation bias was calculated as the difference between the estimated and generating value. Figure S1 shows estimation bias for each group-level parameter.

**Figure S-1.**
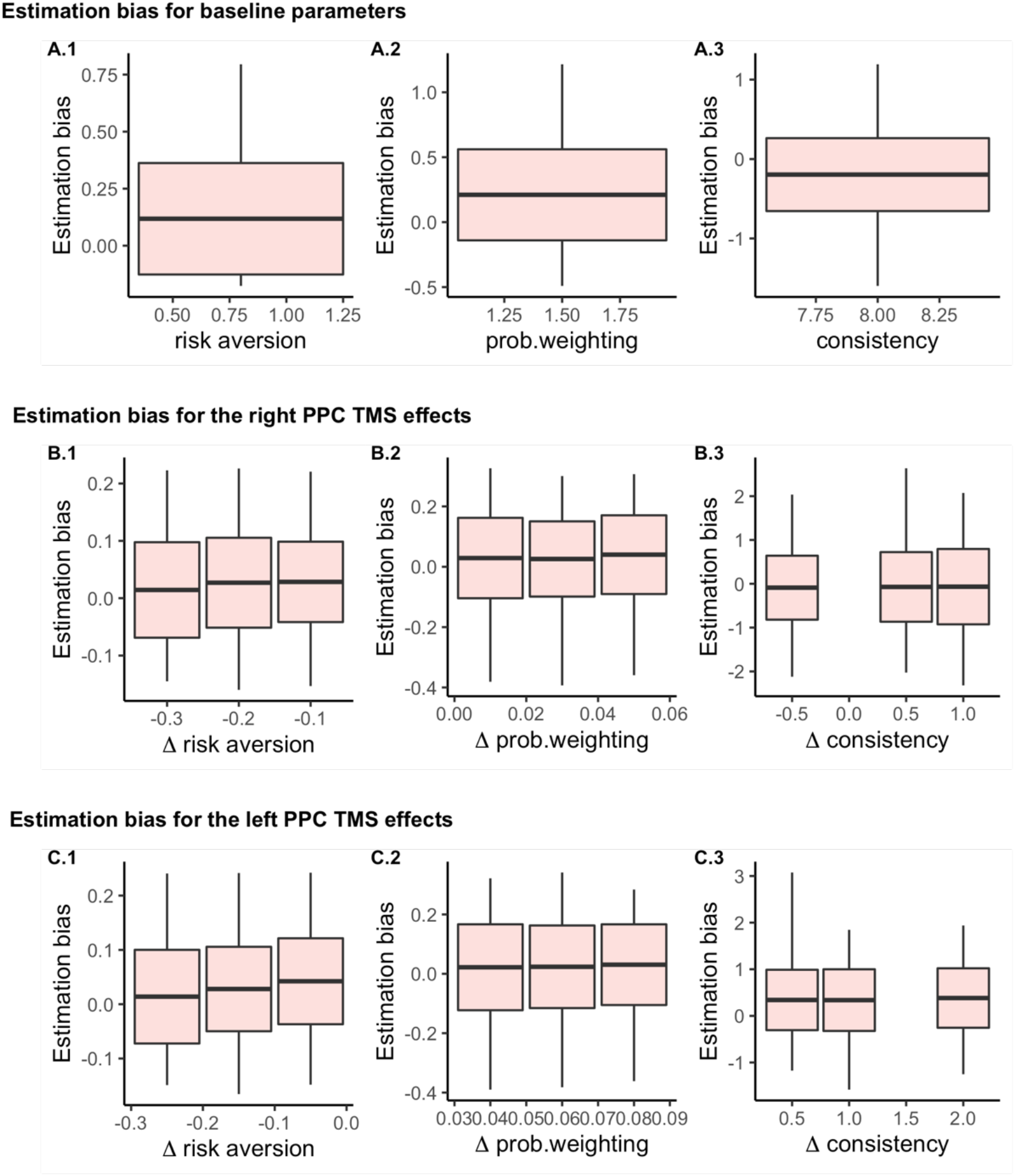
Estimation bias as the difference between recovered and generating parameter values. Each boxplot represents mean±SD, as well as min and max for each parameter estimation bias.

Additionally, we calculated the recovery rate for each parameter, i.e., the percent of cases where the generating parameter value belonged to the 95% HDI. The obtained recovery rates were as follows:: *μ*_*r*^0^_ 0.96, *μ*_*γ*^0^_ 0.92, *μ*_*τ*^0^_ 0.93, *μ*_Δ*r^right^*_ 0.96, *μ*_Δ*γ^right^*_ 0.97, *μ*_Δ*τ^right^*_ 0.97, *μ*_Δ*r^left^*_ 0.97, *μ*_Δ*γ^left^*_ 0.95, *μ*_Δ*τ^left^*_ 0.95. For all parameters measuring the TMS effects the recovery rate was above 95%. Overall, the parameter recovery procedure suggests that all parameters of interest can be recovered well.

### Baseline prospect theory parameter estimates

**Figure S-2.**
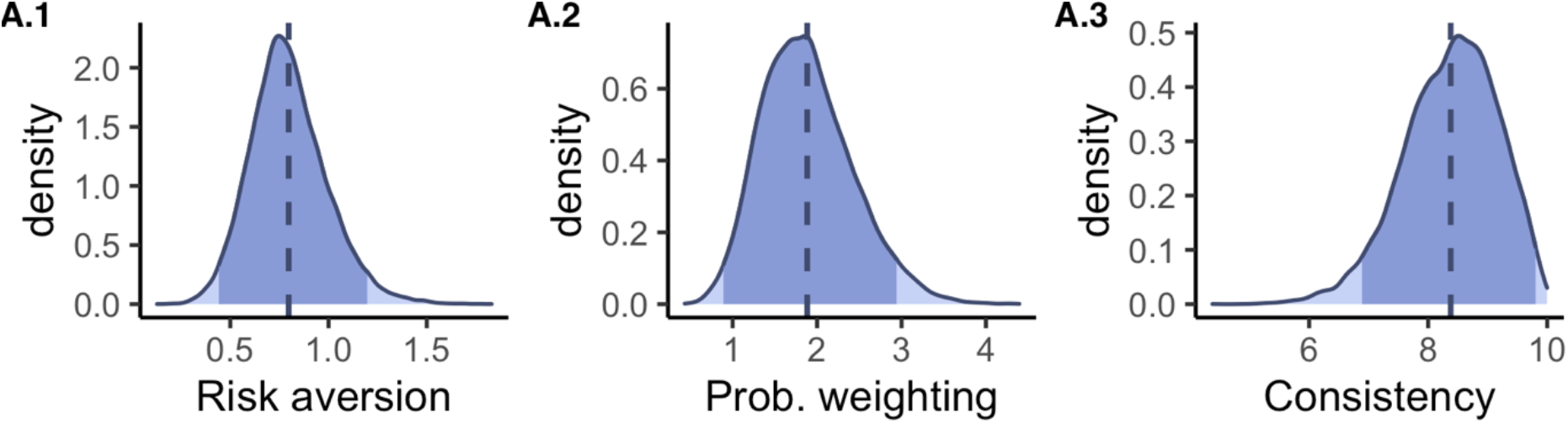
Baseline group-level risk preference parameters: **A.1** Estimated posterior distribution for risk aversion (*μ*_*r*^0^_), **A.2** Estimated posterior distribution for probability weighting (*μ*_*γ*^0^_), **A.3** Estimated posterior distribution for consistency (*μ_τ^0^_*). The shaded area under the curve corresponds to 95% HDI. Dashed vertical line indicates the mean (point estimate) of the posterior distribution.

